# Prediction of compressive strength of vertebral body with metastatic lesions based on quantitative computed tomography-based subject-specific finite element models

**DOI:** 10.64898/2026.03.03.709247

**Authors:** Rajdeep Ghosh, Eddie Shearman, Robert Roger, Marco Palanca, Enrico Dall’Ara, Damien Lacroix

## Abstract

Pathologic vertebral fractures are a major complication in metastatic spine disease. However, current clinical scores, such as Spinal Instability Neoplastic Score (SINS), show limited predictive capability, particularly within the indeterminate range where most clinical uncertainty lies. This study aimed to develop and evaluate quantitative computed tomography (qCT)-based subject-specific finite element (SSFE) models to predict vertebral strength in presence of different metastatic lesion types. Twelve *ex vivo* human spine segments, each containing one metastatic (n=12) and one adjacent control vertebra (n=12), were scanned using qCT and calibrated using a calibration phantom. Homogenised nonlinear finite element models were developed with spatially heterogeneous, isotropic, density-dependent material properties and loaded under uniaxial compression corresponding to 1.9% apparent strain. Ultimate failure load, stiffness, and strain distributions were compared between metastatic and control vertebrae. Predicted failure load ranged from 0.2 kN to 6.2 kN (mean. ± standard deviation: 1.8 ± 1.6 kN metastatic; 1.7 ± 1.5 kN control), with no statistically significant difference between groups (*p* > 0.05). Normalised failure load varied widely, reflecting lesion-specific mechanical heterogeneity. Lytic lesions generally weakened vertebrae, whereas mixed and blastic lesions occasionally enhanced strength, likely due to localised sclerosis or reactive bone formation. High compressive axial strains (greater than 0.019) were frequently concentrated near the endplates, particularly in lytic vertebrae. qCT-derived bone mineral density strongly correlated with failure load (R² = 0.74–0.77). These findings highlight the complexity of metastatic vertebral mechanics and demonstrate that qCT-based SSFE modelling provides a quantitative framework for assessing fracture risk, complementing conventional imaging-based tools.

## 1. Introduction

Cancer comprises a heterogeneous group of diseases (Brown et al. 2023) characterised by uncontrolled cell proliferation and metastasis. It remains a major global health burden and is the second leading cause of mortality worldwide (Siegel et al. 2022). The presence of secondary bone metastases, with over 70% of bone metastases occurring in the spine (Cronin et al. 2018; Sutcliffe et al. 2013), compromises the mechanical competence of vertebrae (Anderson et al. 2026) by altering their microarchitecture and bone mineral density (BMD) distribution. This degradation may occur through a reduction in BMD, as seen in osteolytic metastases; an increase in local BMD and mineralisation, characteristic of blastic or osteosclerotic metastases; or a combination of both effects, as observed in mixed metastases (Cavazzoni et al. 2023). To identify vertebrae at high risk of pathological fracture, scoring systems such as the Spinal Instability Neoplastic Score (SINS) have been proposed (Fourney et al., 2011). However, the SINS has limitations, particularly in its moderate specificity (79.5%) (Fisher et al., 2014), which may result in overtreatment of patients already undergoing aggressive cancer therapies, and in a large number of cases classified as indeterminate (i.e. no recommendation provided by the SINS). Accordingly, there is an unmet need for reliable predictive tools that can assist clinicians in assessing vertebral failure risk in patients with spinal metastases and guide timely, targeted treatment decisions.

Subject-specific finite element (SSFE) models derived from quantitative computed tomography (qCT) scans have shown promising accuracy in estimating vertebral stiffness and failure loads *ex-vivo* (R² = 0.82 for stiffness; R² = 0.78–0.86 for ultimate load) (Wang et al., 2012; Dall’Ara et al., 2012; Crawford et al., 2003). However, the predictive accuracy of failure loads in such SSFE models was found to be limited in mechanically induced lesions (using 0.5% solution agarose gel) in otherwise healthy vertebrae (Groenen et al. 2018). On the other hand, SSFE models developed from high-resolution CT scans have been shown to be accurate in predicting the failure in vertebral slices with different types of metastatic lesions (Gibson et al. 2025; Soltani et al. 2024; Stadelmann et al. 2020; Vialle et al. 2015). Costa et al. (2019) addressed the challenge of assessing vertebral fracture risk in patients with lytic metastases by developing an SSFE approach based on qCT. The study analysed thoracolumbar spine CT scans from eight patients presenting vertebral lytic metastases with SINS in the indeterminate unstable range (7–12). For each patient, one or two vertebrae with lytic lesions were modelled alongside adjacent vertebrae without radiologically visible lesions, which served as internal controls. Interestingly, the authors observed that a comparable number of vertebrae with lytic lesions exhibited either higher or lower mechanical properties than their controls, with only about half of the patients demonstrating significantly weaker metastatic vertebrae. These findings suggest that lytic lesions do not uniformly compromise vertebral strength, underscoring the complexity of fracture risk assessment and the limitations of clinical scoring systems such as SINS. While this study provided an important computational framework, it focused solely on lytic metastases and a relatively small patient cohort, without incorporating blastic or mixed lesion types, which are also clinically relevant. In a comparative study, Stadelmann et al. (2020) evaluated the impact of various metastatic lesion types—lytic, mixed, and blastic—on vertebral strength prediction using SSFE and micro-FE models. The results demonstrated that SSFE models provide an efficient and accurate approach for estimating failure risk in vertebrae with secondary metastatic involvement. Roger et al. (2025) investigated the intra- and inter-operator reproducibility of a qCT-based SSFE modelling pipeline for predicting vertebral strength in vertebrae with lytic lesions. The authors reported high intra-operator consistency quantified by percentage of precision error (ultimate force: 1.5%; ultimate strength: 1.3%), with only modest variability observed in the predicted mechanical properties across operators (ultimate force: 3.6%; ultimate strength: 2.5%). However, to the best of the authors’ knowledge, no prior studies have systematically investigated the effect of different types of metastatic lesions on vertebral strength by directly comparing individual metastatic vertebrae with their adjacent radiological controls.

This study aims to develop SSFE models from qCT scans of cadaveric spine segments to investigate the effect of metastatic involvement on vertebral strength using within-subject comparisons. Particular emphasis is placed on lytic and mixed lesions, with an exploratory assessment of a blastic lesion, to evaluate how lesion type influences failure risk relative to adjacent radiological controls. The proposed comparative computational approach allows assessment of whether vertebrae with different lesion types should be considered at higher risk compared to their adjacent, lesion-free counterparts. Considering that each metastatic lesion type has distinct tissue properties, these SSFE models enable capturing their differential effects on vertebral failure load. This is particularly important when comparing individual metastatic vertebrae with adjacent, lesion-free controls within the same spinal segment, allowing a more comprehensive biomechanical assessment.

## 2. Materials and Methods

### 2.1 Medical imaging and segmentation

qCT scans of 12 cadaveric spine segments, each comprising four vertebrae, were obtained and scanned using a qCT scanner (Aquilion ONE, Toshiba, Japan) following an optimised bone imaging protocol (tube current: 200 mA; tube voltage: 120 kVp; slice thickness: 1 mm; maximum in-plane resolution: approximately 0.45 mm). The same imaging parameters were used to separately scan an offline European Spine Phantom (Quality Assurance in Radiology and Medicine, PTW Freiburg GmbH, Germany), which was later used to calibrate the qCT scans of these spine segments.

Each spine segment included one vertebra with one or more metastatic lesions, one adjacent control vertebra without radiological evidence of metastasis, and two additional cranial and caudal vertebrae used for fixation using polymethylmethacrylate (PMMA). The metastatic lesions were classified based on scans from high-resolution µCT imaging (isotropic voxel size 39 μm; current 114 mA; voltage 70kVp; integration time 300 ms; power 8W; VivaCT80, Scanco, Switzerland). Table 1 shows the details of the donors, levels included in the spine segments, modelled vertebral levels and their condition (metastatic vs control). Each metastatic and control vertebrae, excluding the posterior elements, were then manually segmented in the 3D Slicer image processing platform (3D Slicer, v5.6.2) using a suitable level of smoothing to remove unwanted noise from the segmentations.

**Table 1:**
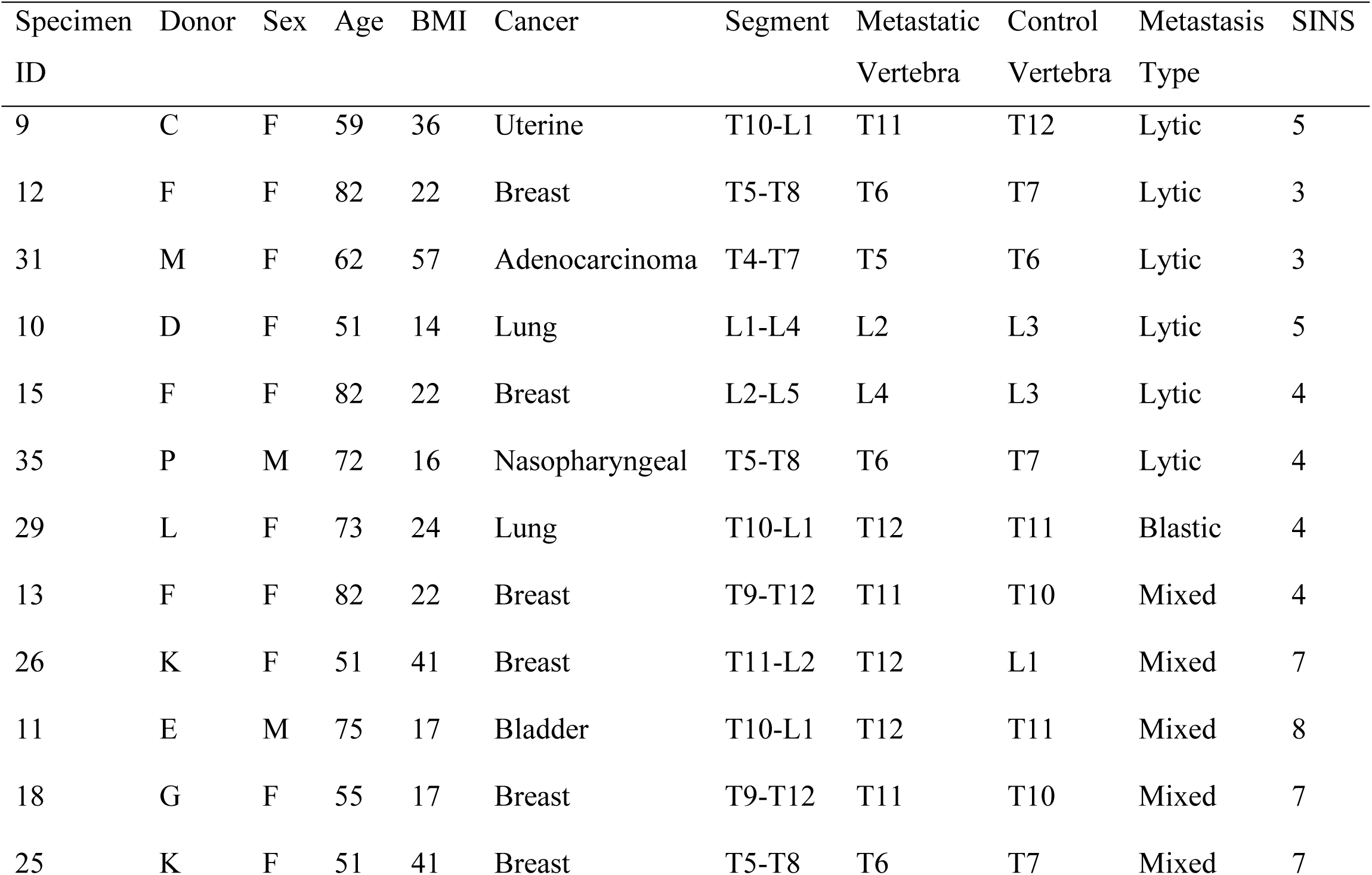
Details of the donor, levels included in the spine segments, modelled vertebral levels and their condition based on microCT images and the radiographic SINS (Palanca et al. 2021)

### 2.3 Identification of endplates and anatomical alignment

The segmented vertebrae in each spine segment were then exported to ANSYS Spaceclaim (Ansys^®^ 2023R2, Ansys Inc., USA) to manually identify the bony endplates at the cranial and caudal surfaces of the vertebral body. The vertebra was further aligned with a global reference system in ANSYS 2023R2, based on an earlier study which reported a reproducible reference system for cadaveric testing of human vertebrae (Danesi et al., 2014). Briefly, two anatomical planes were defined at the cranial and caudal endplates of each vertebra by marking three virtual landmarks in each plane, viz., one at the most anterior end and the other two at the left and right corners in the most posterior end. A mid-plane bisecting these two planes was then created, passing through the centre of the vertebral body. The angle between this mid-plane and the transverse anatomical plane was used to align the vertebra with respect to the global reference frame. Alignment was achieved by defining a new local coordinate system in ANSYS Workbench (2023R2), with its origin positioned at the centroid of the superior endplate and oriented perpendicular to the endplates. The model was then aligned by applying rotations about the axial, anterior–posterior and left–right axes based on the angles between the transverse and bisector planes.

The cross-sectional area *A_m_* of each vertebra was calculated as the area of the vertebral surface at the mid-plane passing through the vertebral body. The minimum height H_min_ of the vertebral body was estimated as the axial distance between the most concave points on each of the vertebral endplates.

### 2.4 Development of Finite Element Models

Each vertebral body geometry was meshed using 10-noded quadratic tetrahedral finite elements with a maximum edge size of 1.0 mm, according to the results of a previous mesh convergence study (Costa et al. 2019; Gibson et al. 2025). Bone was modelled as a heterogeneous, isotropic, bilinear elasto–plastic material. Heterogeneous linear elastic properties were assigned using Bonemat (rolling version 0.2; https://ior-bic.github.io/software/bonemat/index.html), based on BMD values obtained from qCT images that were densitometrically calibrated (Carpenter et al. 2014; Pickhardt et al. 2013) using the European Spine Phantom scanned offline. The resulting calibration curve, shown in Supplementary Figure 1 was used to convert voxel-wise HU values from the qCT images into corresponding qCT-equivalent BMD (ρ_qCT_) values for the vertebral body, including both cortical and trabecular bone, while excluding the posterior elements. These BMD values were subsequently mapped to apparent density ρ_app_ (Les et al. 1994; Schileo et al. 2008) and then into elastic modulus E (Morgan et al. 2003) values using an established density–elasticity relationship [Eq. (1) and Eq. (2)].

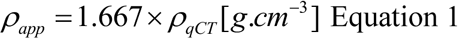

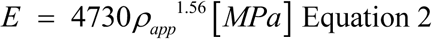

Bone Mineral Content (BMC) was further derived from BMD (in g.cm^-3^) and bone volume (in cm³) using the relations [Eq. (3)] [Eq. (4)]:

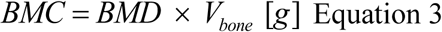

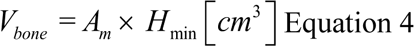

Bone plasticity was modelled using an isotropic von Mises yield criterion based on a density–strength relationship [Eq. (5)] (Morgan and Keaveny, 2001). Bone exhibits tension–compression asymmetry, with tensile yield strengths typically lower than compressive yield strengths (Morgan and Keaveny, 2001). The tensile density–based yield law was used to provide a conservative failure threshold; this ensures that the model does not overpredict the structural capacity. An isotropic hardening rule was applied, incorporating a 95% reduction in the post-yield modulus [Eq. (6)] (Morgan et al., 2003; Bayraktar et al., 2004; Morgan and Keaveny, 2001).

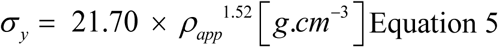

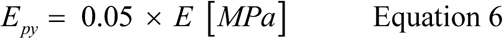

### 2.3 Boundary conditions

The FE models were subjected to a compressive loading condition to simulate physiological axial loading of the vertebral body (Nazarian et al. 2008). An axial displacement was applied to the surface nodes of the most cranial endplate, enforcing a uniform compression corresponding to 1.9% of the vertebral height H_min_, which was adopted as the failure criterion (Wang et al. 2012; Keaveny et al. 2014) to determine the strength of the vertebra. This displacement-controlled approach allows for consistent comparison of vertebral strength across specimens, independent of geometry or material heterogeneity. The nodes on the caudal endplate were fully constrained in all translational and rotational degrees of freedom to replicate a fixed boundary condition and ensure model stability during loading. In addition, the loading direction, defined in the anatomically aligned local coordinate system, ensured physiologically realistic stress and strain distributions within both cortical and trabecular compartments of the vertebral body. Each finite element model comprised approximately 3 million degrees of freedom and required about 2 hours to solve in ANSYS 2023R2 (Ansys Inc., USA). Simulations were performed on Stanage, the University of Sheffield’s high-performance computing cluster, using parallel distributed memory across up to 32 cores based on one node of dual 32-core Intel Xeon Platinum 8358 processors with 256 GB RAM.

### 2.4 Determination of biomechanical metrics

To quantify the structural strength of the vertebral body under uniaxial compression, the ultimate force, or failure load (F_L_), for each tested vertebra was determined. It was defined as the sum of the reaction forces at the inferior endplate when the applied axial displacement (Δ*l*) reached 1.9% of apparent strain (Wang et al. 2012; Keaveny et al. 2014; Costa et al. 2019). Apparent strain (*ε_app_*) was calculated as the ratio of axial displacement (Δ*l*) to the minimum height of the vertebra (*H_min_*) [Eq. (7)]. In addition, load-displacement and stress-strain curves were plotted for each vertebra. Ultimate stress (*σ_ult_*) was defined as the ratio of failure load to the cross-sectional area at the mid-transverse plane (*A_m_*) [Eq. (8)].

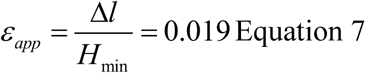

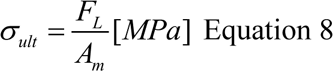

A normalised failure load (*F_norm_*), was defined as the ratio between the failure load of the metastatic vertebra (*F_ult-m_*) and that of the adjacent control (*F_ult-c_*) one [Eq. (9)]. *F_norm_* is a dimensionless parameter, calculated to minimise the variation in intra-patient bone size, geometry and density. Structural stiffness (*K_g_*) was calculated as the slope of the linear range of the load-displacement curves [Eq. (10)]. Apparent stiffness (*K_app_*) was calculated as the normalisation of the structural stiffness with the ratio of the minimum height of the vertebral body to the cross-sectional area of the vertebral body at the mid-transverse plane [Eq. (11)]. Total minimum principal strain contour plots of the tested specimens were plotted and analysed to understand how different lesions affect the load distribution within each vertebra.

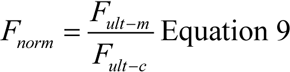

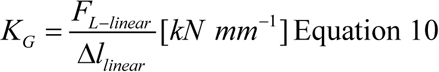

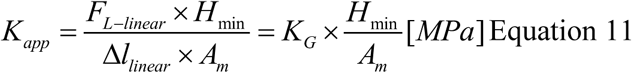

### 2.5 Statistical analysis

Statistical analysis was performed to compare FE-predicted mechanical metrics between metastatic vertebrae and their adjacent radiological control vertebrae (n = 12 pairs). Normality of the data was assessed using the Shapiro–Wilk test. As normality could not be assumed for all variables, non-parametric statistical tests were employed. Differences in failure load and compressive strength between control and metastatic vertebrae were evaluated using the Mann–Whitney U test. Statistical significance was defined at *p* < 0.05. As the median is less affected by skewness and outliers, the median value of the total minimum principal strain was used to represent the characteristic strain magnitude in each vertebra.

Linear regression analyses were performed to assess the relationships between FE-predicted failure load and compressive strength with qCT-derived BMD and BMC in control, metastatic, and pooled vertebrae. The strength of association was quantified using the coefficient of determination (R²).

## 3 Results

The geometrical and densitometric properties of each tested vertebral body are reported in Table 2. Mean qCT equivalent BMD over the pooled data ranged between 0.07 g.cm^-3^ to 0.40 g.cm^-3^ (mean and standard deviation of 0.19±0.08 g cm^-3^ for metastatic; 0.18±0.09 g cm^-3^ for controls). The mean BMC over the pooled data ranged between 1.39 g to 8.93 g (mean and standard deviation of 3.04±1.55 g for metastatic; 3.08±2.14 g for controls).

**Table 2:** Details of the measured minimum height of vertebra, cross-sectional area, qCT equivalent BMD, BMC and lesion size.

| Specimen ID | Segment | Level | Condition | H<sub>min</sub> (mm) | A<sub>m</sub> (mm<sup>2</sup>) | qCT equivalent BMD (g cm<sup>-3</sup>) | BMC (g) | Lesion Size (VB%(cm<sup>3</sup>)) |
| --- | --- | --- | --- | --- | --- | --- | --- | --- |
| 9 | T10-L1 | T11 | Lytic | 19.2 | 955 | 0.19 | 3.64 | 2 (0.4) |
|  |  | T12 | Control | 21.0 | 1137 | 0.16 | 3.93 |  |
| 12 | T5-T8 | T6 | Lytic | 21.2 | 455 | 0.17 | 1.67 | 5 (0.6) |
|  |  | T7 | Control | 17.9 | 510 | 0.16 | 1.53 |  |
| 31 | T4-T7 | T5 | Lytic | 16.0 | 485 | 0.23 | 1.86 | 6 (0.8) |
|  |  | T6 | Control | 21.0 | 533 | 0.23 | 2.86 |  |
| 10 | L1-L4 | L2 | Lytic | 26.0 | 874 | 0.07 | 1.77 | 36 (10.1) |
|  |  | L3 | Control | 26.0 | 955 | 0.07 | 1.88 |  |
| 15 | L2-L5 | L4 | Lytic | 29.0 | 781 | 0.14 | 3.32 | 20 (5.2) |
|  |  | L3 | Control | 26.0 | 798 | 0.15 | 3.12 |  |
| 35 | T5-T8 | T6 | Lytic | 18.0 | 500 | 0.20 | 1.83 | 10 (1.0) |
|  |  | T7 | Control | 18.0 | 465 | 0.16 | 1.40 |  |
| 29 | T10-L1 | T12 | Blastic | 22.8 | 933 | 0.11 | 2.50 | 12 (3.6) |
|  |  | T11 | Control | 22.1 | 956 | 0.02 | 0.55 |  |
| 13 | T9-T12 | T11 | Mixed | 21.0 | 761 | 0.16 | 2.67 | 12 (2.2) |
|  |  | T10 | Control | 23.0 | 676 | 0.17 | 2.73 |  |
| 26 | T11-L2 | T12 | Mixed | 24.8 | 971 | 0.30 | 7.30 | 19 (5.3) |
|  |  | L1 | Control | 27.8 | 978 | 0.32 | 8.93 |  |
| 11 | T10-L1 | T12 | Mixed | 22.0 | 1050 | 0.11 | 2.62 | 4 (1.4) |
|  |  | T11 | Control | 20.8 | 1144 | 0.13 | 3.16 |  |
| 18 | T9-T12 | T11 | Mixed | 22.5 | 752 | 0.19 | 3.29 | 20 (5.0) |
|  |  | T10 | Control | 21.9 | 663 | 0.17 | 2.57 |  |
| 25 | T5-T8 | T6 | Mixed | 16.8 | 559 | 0.40 | 4.05 | 31 (4.3) |
|  |  | T7 | Control | 20.0 | 647 | 0.34 | 4.51 |  |

The predicted load–displacement and stress–strain responses of vertebrae with and without metastatic lesions exhibited substantial variability in their mechanical properties (Figure 2). Failure load for each of the modelled vertebrae compared to adjacent controls in each spine segment are reported in Table 3. The ultimate failure load across the pooled dataset ranged from 0.2 to 6.2 kN, with comparable mean values for metastatic (1.8 ± 1.6 kN) and control vertebrae (1.7 ± 1.5 kN). Most lytic lesions exhibited a normalised failure load below unity, whereas the blastic vertebra showed an increased failure load (Figure 3b). Mixed lesions demonstrated heterogeneous behaviour, with the majority showing reduced normalised failure loads. One lytic specimen (Specimen ID 10) deviated from this trend, exhibiting a relatively high normalised failure load despite a large lesion. Vertebral ultimate stress (Figure 3a), structural stiffness (Figure 3c), and apparent stiffness (Figure 3d) showed overlapping ranges between metastatic and control vertebrae, with no systematic differences observed across the pooled dataset. Summary statistics for all mechanical metrics are reported in Table 4.

**Figure 1.**
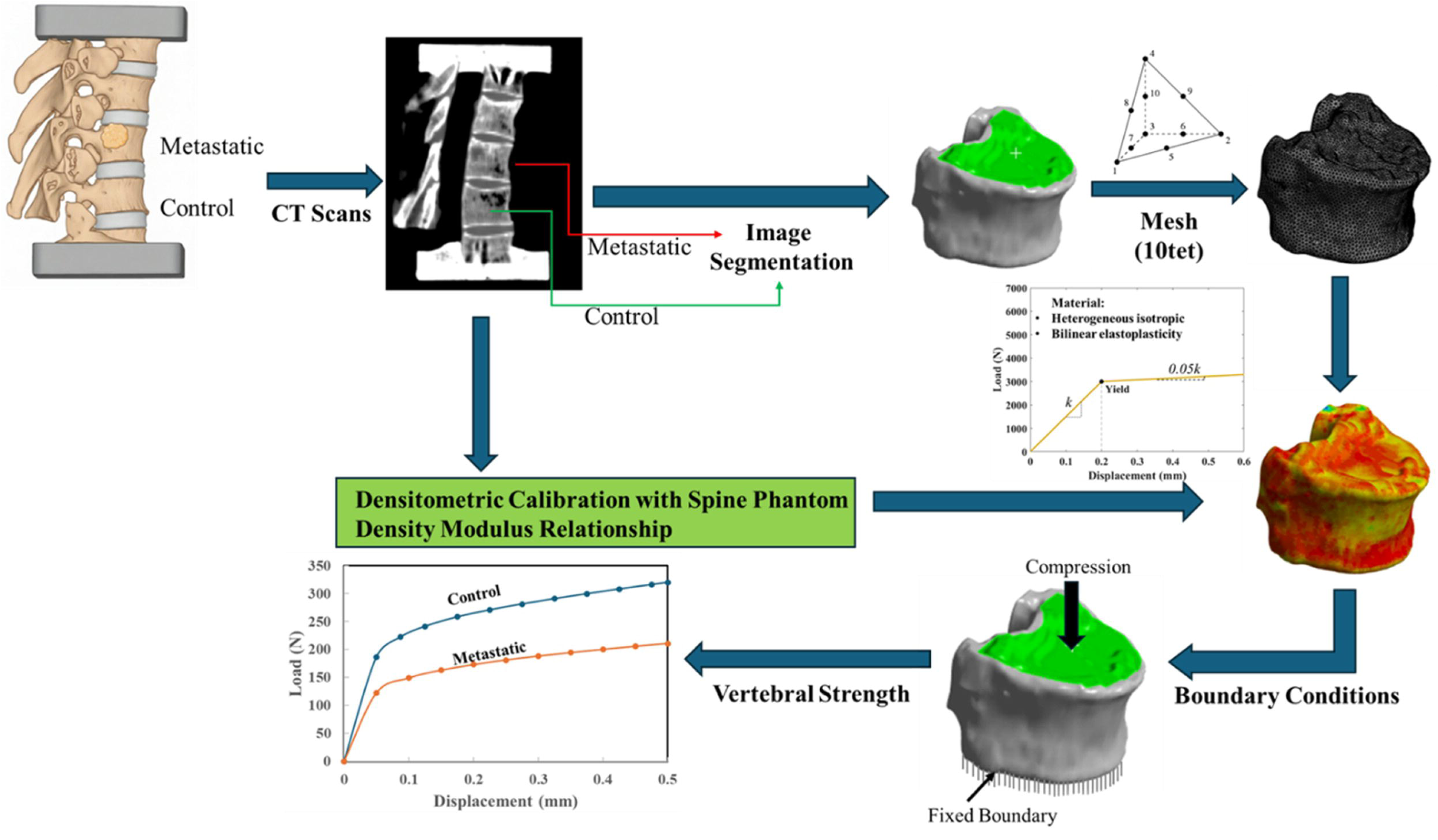
Reproducible computational pipeline from qCT to development of SSFE models to predict biomechanical metrics such as failure load, stress, strain etc.

**Figure 2.**
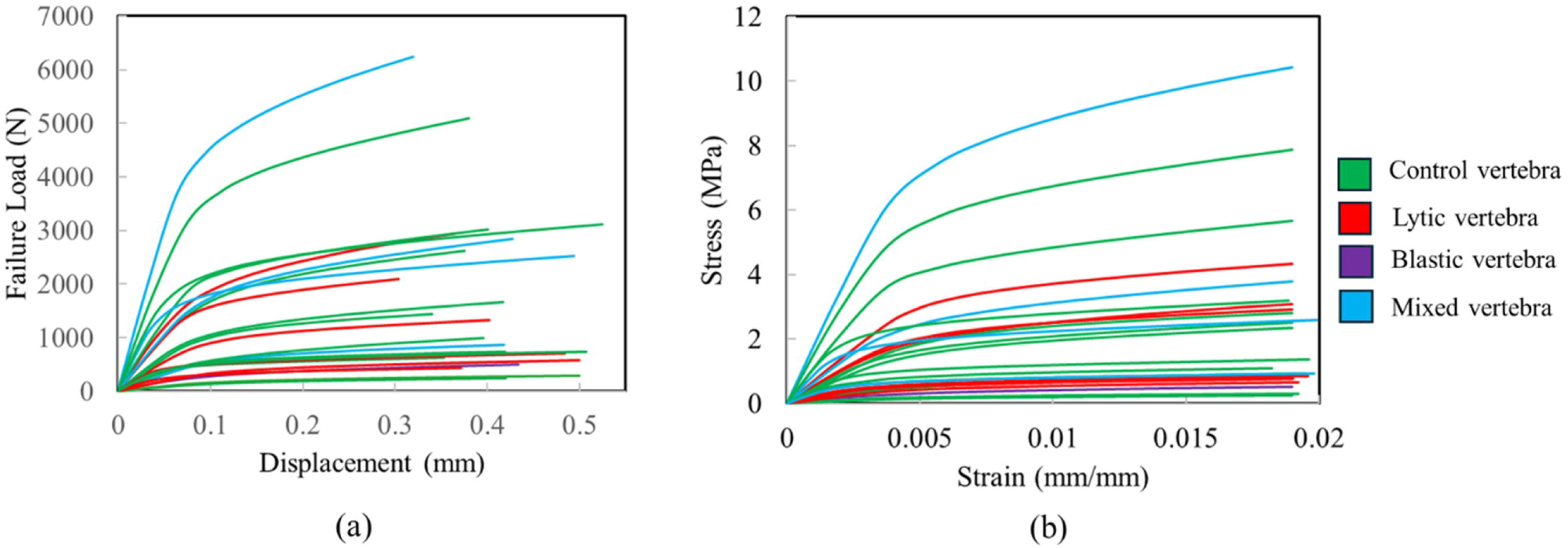
Load-displacement and stress-strain curves for metastatic vertebrae (lytic, mixed, and blastic) and their adjacent radiological control vertebrae.

**Figure 3.**
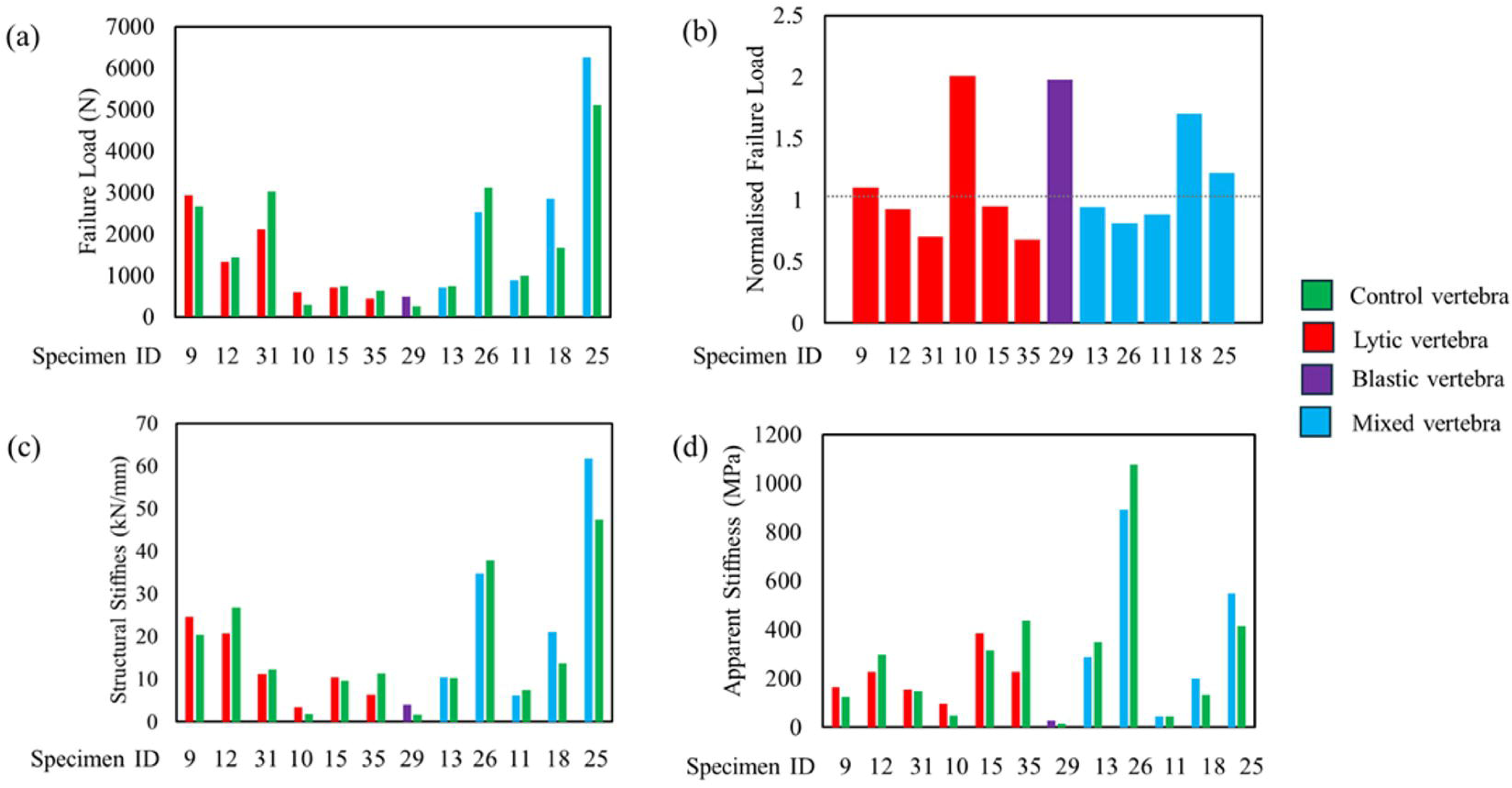
(a) Ultimate Failure Load; (b) Normalised failure load; (c) structural stiffness; and (d) apparent stiffness predicted for metastatic vertebrae (lytic, mixed, and blastic) and their adjacent radiological control vertebrae.

**Table 3:** Details of the median total minimum principal strain, ultimate failure load, percentage difference in failure load between metastatic and adjacent control and endplates experienced high compressive strains (>0.019)

| Specimen ID | Segment | Level | Condition | Median Total Minimum Principal Strain ($\mu\epsilon$) | F<sub>L</sub> (N) | Superior Endplate | Inferior Endplate |
| --- | --- | --- | --- | --- | --- | --- | --- |
| 9 | T10-L1 | T11 | Lytic | 9540 | 2940 | ✓ |  |
|  |  | T12 | Control | 9846 | 2667 |  |  |
| 12 | T5-T8 | T6 | Lytic | 13460 | 1326 | ✓ |  |
|  |  | T7 | Control | 11840 | 1436 |  |  |
| 31 | T4-T7 | T5 | Lytic | 9120 | 2112 | ✓ |  |
|  |  | T6 | Control | 14212 | 3014 | ✓ |  |
| 10 | L1-L4 | L2 | Lytic | 8910 | 586 | ✓ |  |
|  |  | L3 | Control | 5320 | 290 |  |  |
| 15 | L2-L5 | L4 | Lytic | 7910 | 703 | ✓ |  |
|  |  | L3 | Control | 6980 | 739 | ✓ |  |
| 35 | T5-T8 | T6 | Lytic | 4920 | 431 | ✓ |  |
|  |  | T7 | Control | 6390 | 633 |  |  |
| 29 | T10-L1 | T12 | Blastic | 4170 | 494 |  |  |
|  |  | T11 | Control | 7610 | 249 |  |  |
| 13 | T9-T12 | T11 | Mixed | 6150 | 699 |  |  |
|  |  | T10 | Control | 7630 | 744 | ✓ | ✓ |
| 26 | T11-L2 | T12 | Mixed | 7600 | 2528 | ✓ | ✓ |
|  |  | L1 | Control | 8800 | 3111 | ✓ |  |
| 11 | T10-L1 | T12 | Mixed | 5900 | 874 |  |  |
|  |  | T11 | Control | 3970 | 992 |  |  |
| 18 | T9-T12 | T11 | Mixed | 8760 | 2840 |  | ✓ |
|  |  | T10 | Control | 6850 | 1668 |  |  |
| 25 | T5-T8 | T6 | Mixed | 10520 | 6247 |  | ✓ |
|  |  | T7 | Control | 12080 | 5100 | ✓ |  |

**Table 4:** Summary statistics of FE-predicted mechanical metrics for control and metastatic vertebrae.

| **Mechanical Metric** | **Group** | **Mean±SD** | **Range** |
| --- | --- | --- | --- |
| ***Ultimate Failure Load (kN)*** | Control | 1.7±1.5 | 0.2-6.2 |
|  | Metastatic | 1.8±1.6 |  |
| ***Ultimate Stress (MPa)*** | Control | 2.4±2.3 | 0.3-10.4 |
|  | Metastatic | 2.6±2.8 |  |
| ***Structural Stiffness (kN/mm)*** | Control | 16.7±14.1 | 1.6-61.8 |
|  | Metastatic | 17.8±16.8 |  |
| ***Apparent Stiffness (MPa)*** | Control | 283.5±290.3 | 13.7-1076.8 |
|  | Metastatic | 270.4±242.9 |  |

Mann–Whitney U test revealed no statistically significant difference in failure load between control and metastatic vertebrae (U = 67, *p* = 0.885). Similarly, Mann–Whitney U test showed no statistically significant difference in compressive strength (U = 71, *p* = 0.977).

Figure 4 illustrates the relationship between the predicted failure load with mean qCT-equivalent BMD and mean BMC for control and metastatic vertebrae. A good positive correlation was observed between BMD and failure load, with R² values of 0.77, 0.74, and 0.75 for control, metastatic, and pooled data, respectively. When comparing the BMD and the ultimate strength of the individual tested vertebrae, a moderate correlation was observed, with R² values of 0.55, 0.68, and 0.61 for the control, metastatic, and pooled data, respectively. BMC showed a weak association with failure load, with corresponding R² values of 0.23, 0.27, and 0.24 for the same groups.

**Figure 4.**
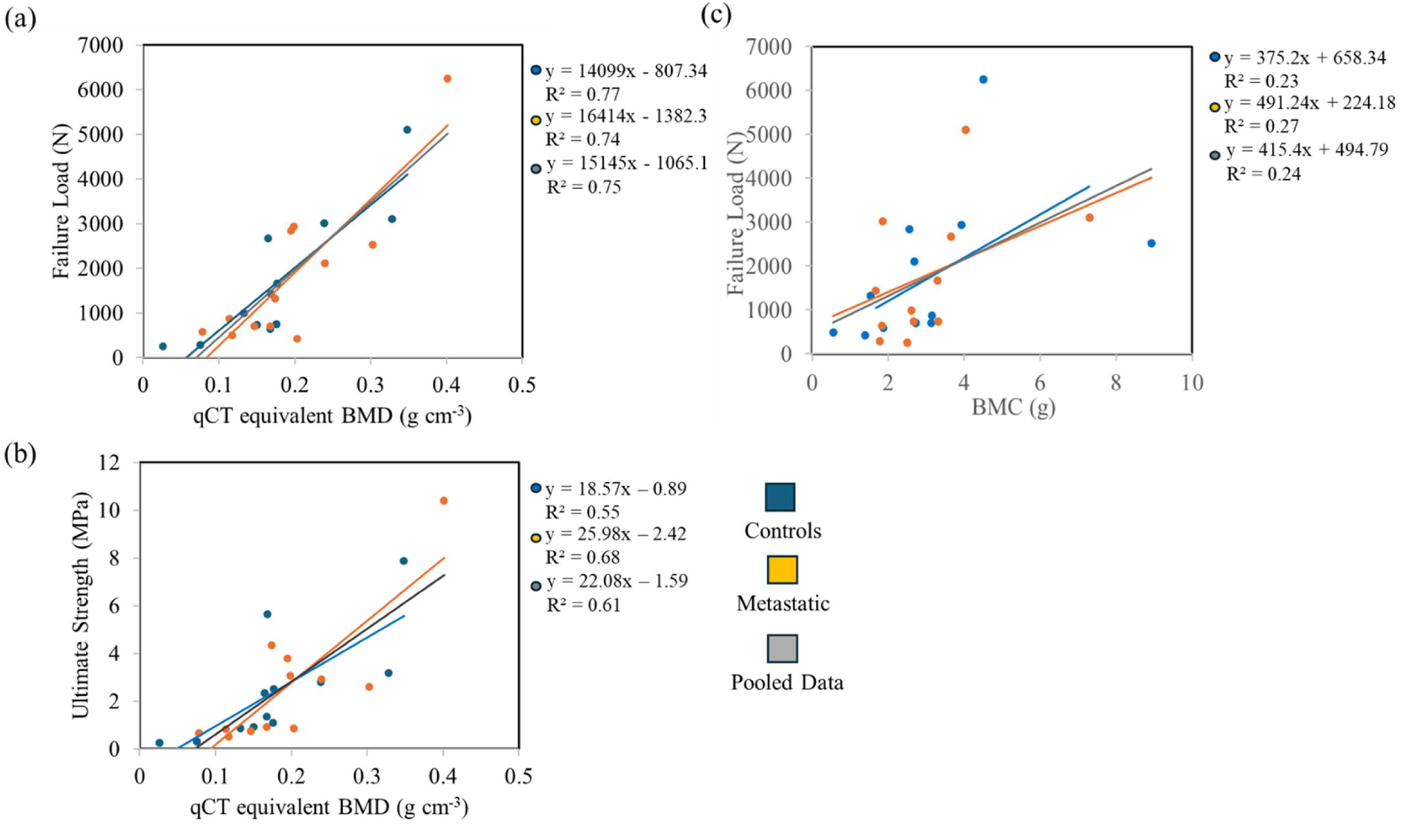
Relationship between qCT equivalent BMD and (a) ultimate failure load predicted; (b) Ultimate Strength; and (c) between BMC (Bone Mineral Content) and ultimate failure load predicted for each of the tested vertebrae.

Frequency plots for each modelled vertebra are presented in Figures 5-6. The strain distributions were skewed in all vertebrae, indicating a non-uniform strain response within the trabecular and cortical regions. Strain (Table 3) across the pooled data ranged between 3970 µε to 14214 µε. However, when summarised at the group level, the mean ± standard deviation of the vertebra-specific median strains was 8081 ± 2580 µε for metastatic vertebrae and 8461 ± 3017 µε for controls. Figures 6-8 also show the strain distribution in the metastatic, and their adjacent control vertebrae for different types of metastatic lesions. Strain distribution pattern shows that higher strains were located primarily at the trabecular bone away from the cortical shell, irrespective of the condition or the type of vertebra. Both controls and metastatic vertebrae were found to be subjected to locally higher strains inside the vertebra than the apparent failure strain of 0.019. High compressive strains were also observed near the endplates in several specimens, irrespective of the pathological condition of the vertebra. Vertebrae exhibiting endplate regions subjected to high compressive strains (>0.019) are summarised in Table 3. Among the 24 vertebrae modelled in this study, 14 showed evidence of endplate involvement: 10 exhibited high strains at the superior endplate, 2 at the inferior endplate, and 2 at both endplates. Notably, all vertebrae with lytic lesions (Figure 5) demonstrated regions of elevated strain (>0.019) at their superior endplates. In contrast, inferior endplate involvement was predominantly observed in vertebrae with mixed lesions (Figure 6). The spinal segment containing the blastic lesion (Figure 6) did not exhibit high strain concentrations at either the superior or inferior endplates in either the metastatic or control vertebra.

**Figure 5.**
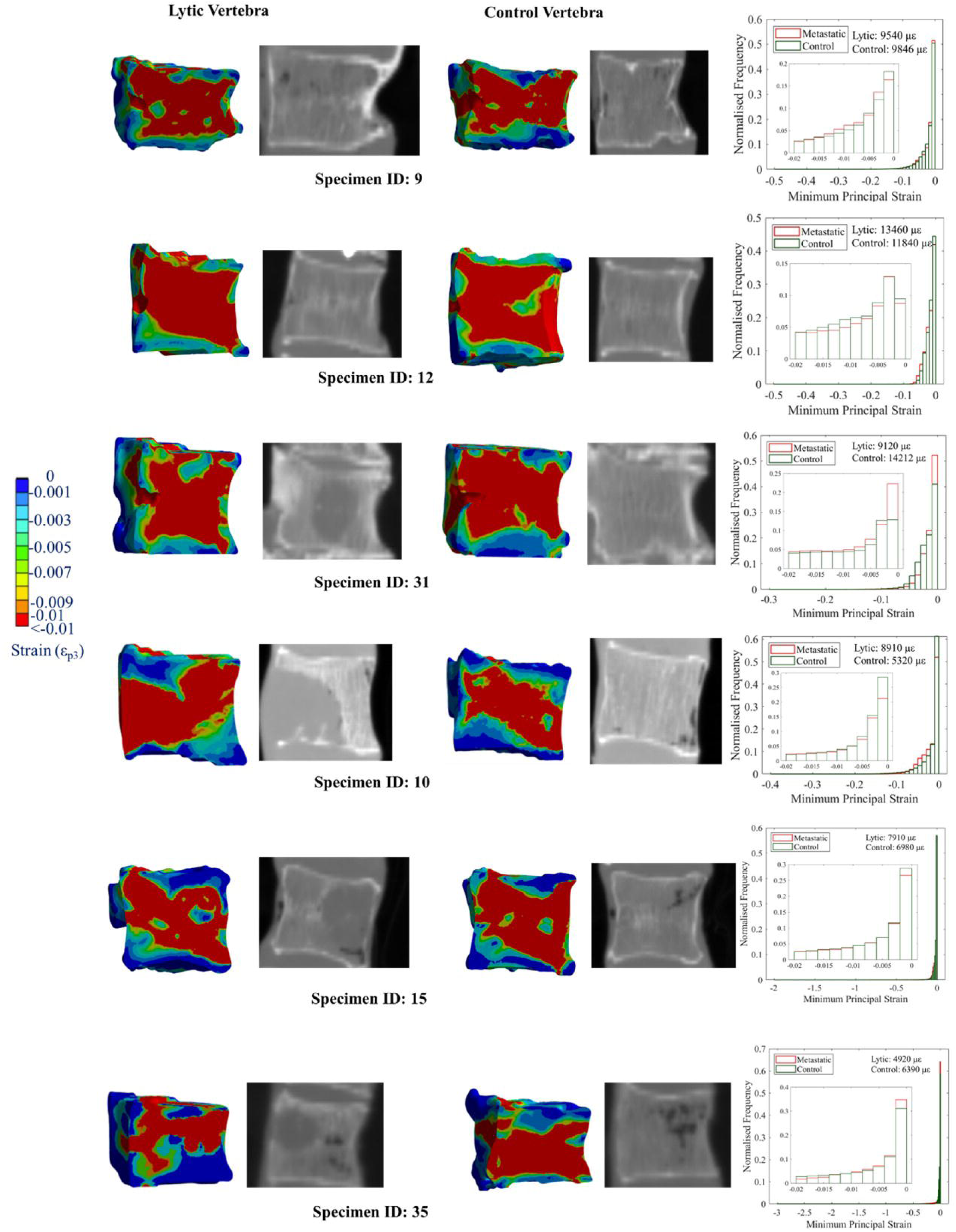
Total minimum principal strain distribution of all the tested lytic cases, along with their adjacent control vertebrae. Strain frequency distribution plots along with reported median strains for each specimen. The zoom in frequency distribution shows strain values that are within failure strain (ε*_app_*=0.019).

**Figure 6.**
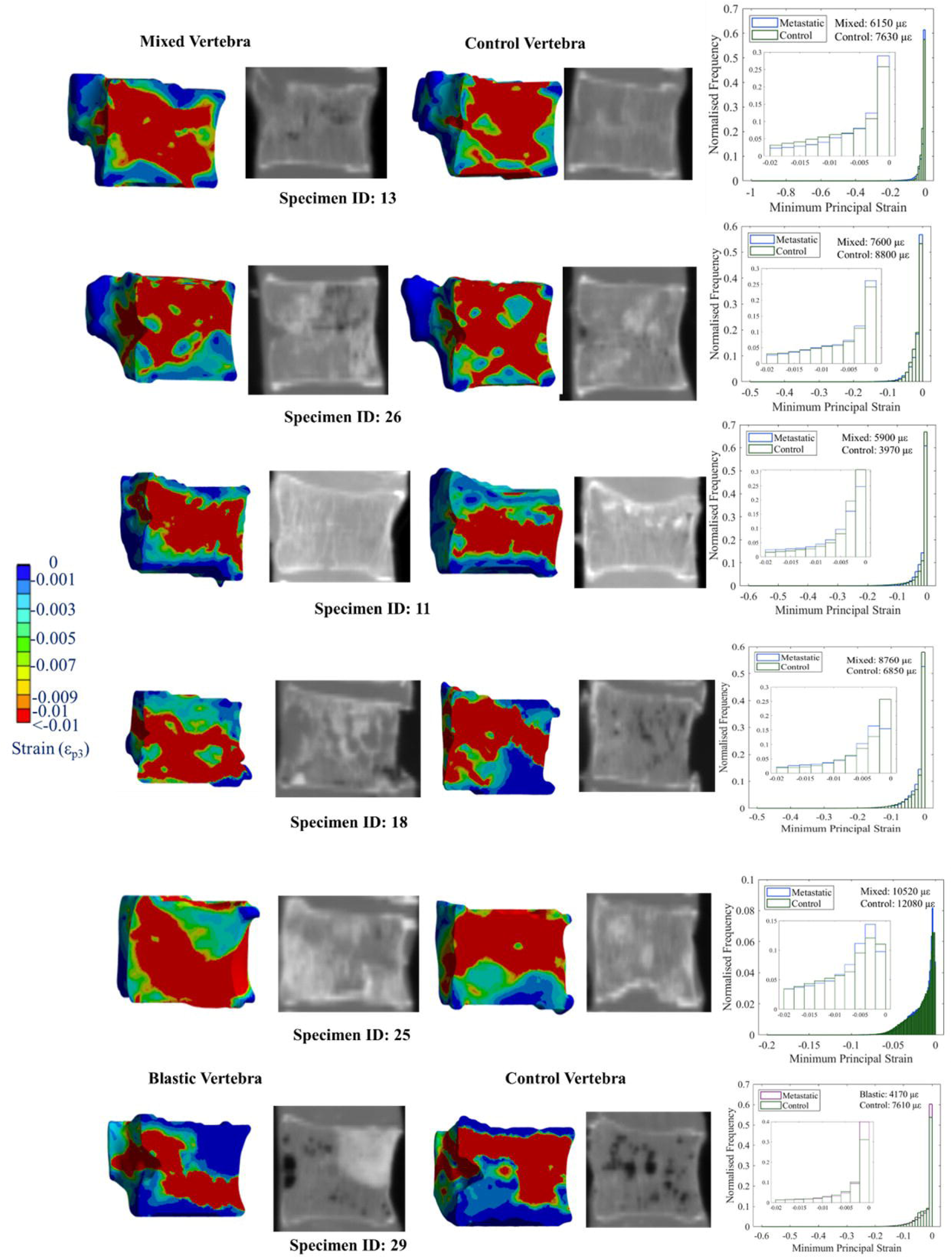
Total minimum principal strain distribution of all the tested mixed and blastic cases, along with their adjacent control vertebrae. Strain frequency distribution plots along with reported median strains for each specimen. The zoom in frequency distribution shows strain values that are within failure strain (ε*_app_*=0.019).

This observation can be further interpreted by examining the spatial distribution of high minimum principal strain regions for each type of metastatic lesion. Figure 7 shows the locations of highly strained regions within lytic, blastic, and mixed metastatic vertebrae. In lytic vertebrae, clusters of elevated compressive strains (exceeding 5%) were predominantly localised around the lytic regions (marked in red in the qCT images), indicating localised loss of load-bearing capacity. In contrast, in the blastic vertebra, highly strained regions (exceeding 7%) were primarily located away from the sclerotic tissue (marked in yellow), within adjacent compliant regions of the vertebral body. For mixed metastatic vertebrae, elevated compressive strains (exceeding 5%) were generally observed near lytic regions and away from blastic tissue. In a subset of specimens, where lytic and blastic tissues were diffusely distributed across the vertebral body, strain localisation was less distinct, making identification of concentrated high-strain regions from the contour plots more challenging.

**Figure 7.**
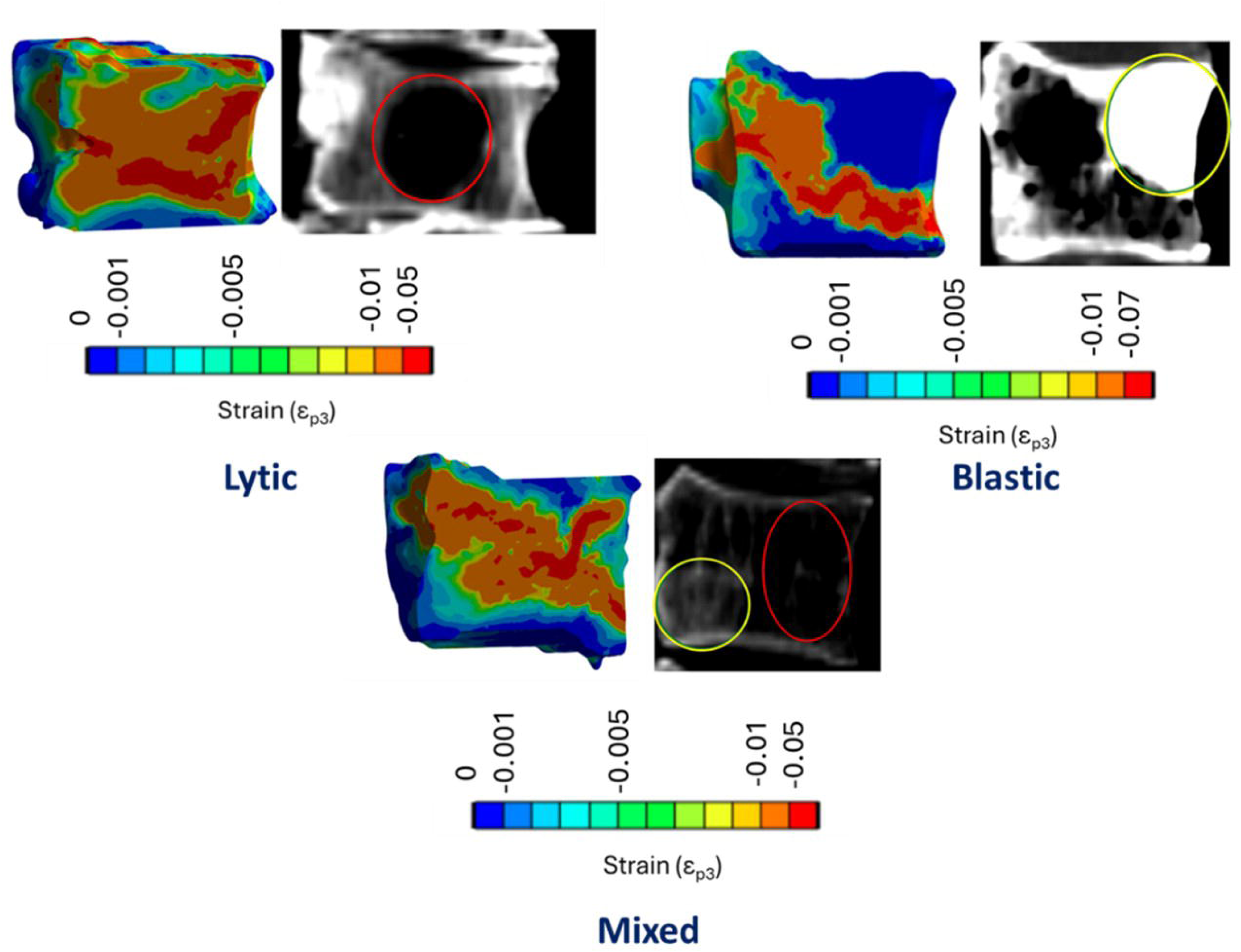
Representative vertebrae illustrating the spatial distribution of high compressive strain regions in different types of metastatic lesions. Strain thresholds shown in the plots are used solely for visualisation of strain localisation and do not represent the failure criterion, which was defined at 1.9% compressive strain.

Equivalent plastic strain was concentrated at the lesion–bone interface in both representative metastatic cases (Figure 8). In specimen ID10 with the large off-centre lytic lesion, plastic deformation was restricted to an interfacial annulus surrounding the lesion cavity and accounted for 2.1% of the vertebral volume, compared with 1.5% in its adjacent control. A similar perilesional localisation was observed in the blastic specimen (ID29), where 1.8% of the vertebral volume entered plasticity, versus 1.9% in the corresponding control. In all cases, plasticity mostly remained confined to the immediate lesion–bone interface, with negligible deformation within the lesion cavity itself or in remote trabecular regions.

**Figure 8.**
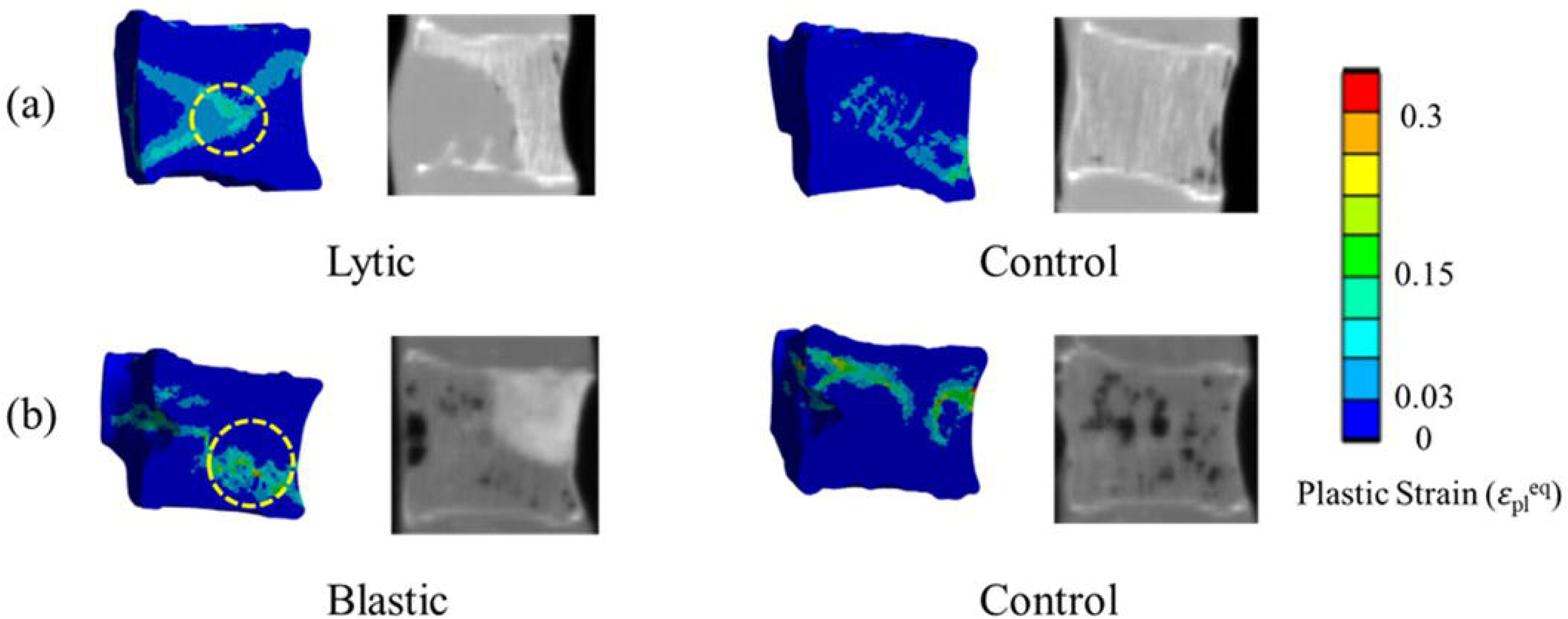
Spatial distribution of equivalent plastic strain in representative lytic and blastic metastatic vertebrae and their adjacent radiological controls. Regions exhibiting high levels of plastic strain are highlighted in yellow, illustrating the extent and localisation of irreversible deformation around metastatic lesions.

Figure 9 shows the relationship between the radiographic SINS and normalised failure load for all metastatic vertebrae. Due to the *ex vivo* nature of the specimens, the pain component of SINS could not be assessed; therefore, the score was based exclusively on radiographic parameters. Within the lower SINS range (0–6), most lytic vertebrae (n = 6) exhibited normalised failure loads below 1, ranging from 0.68 to 1.1. One exception was Specimen ID10, which showed a normalised failure load of 2.01 at a SINS score of 5. Mixed lesions (n = 5) were distributed across both the lower (SINS 4) and potentially higher categories (SINS 7–8), with normalised failure loads ranging from 0.81 to 1.7. Several mixed lesions with SINS scores of 7–8 exhibited normalised failure loads greater than 1. The blastic vertebra (Specimen ID29) showed a high normalised failure load of 1.98 at a SINS score of 4.

**Figure 9.**
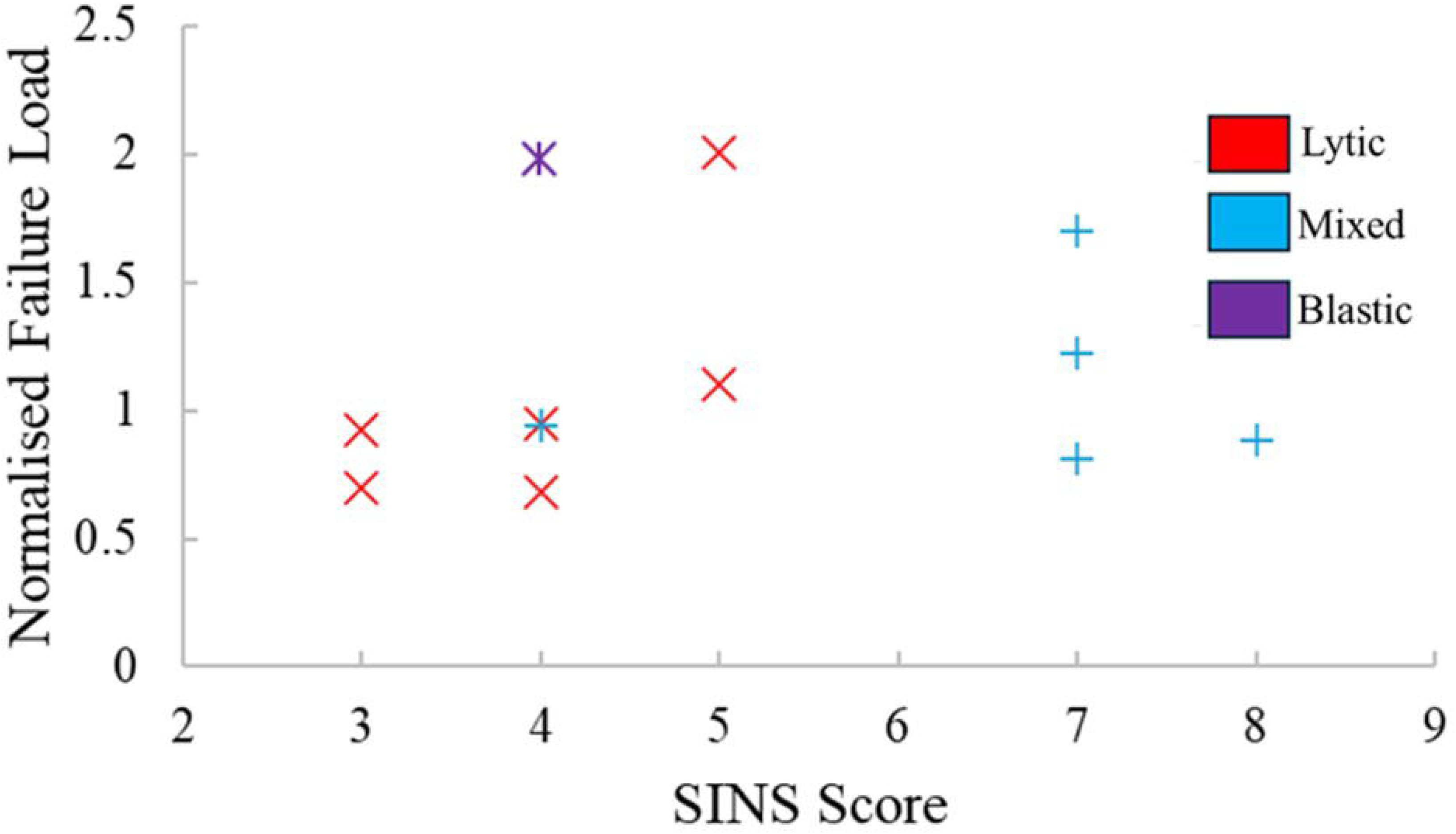
Relationship between radiographic SINS and normalised failure load across all the tested spine segments

## 4 Discussion

This study presented a computational framework to assess vertebral strength in the presence of metastatic lesions using SSFE models derived from qCT imaging. The results demonstrated that ultimate strength, stiffness, and strain distribution vary widely among metastatic vertebrae, reflecting the biological and structural heterogeneity of metastatic involvement. Despite the overall similarity in mean failure load and strength between metastatic and control vertebrae, substantial intra-group variability was observed, underscoring the necessity of personalised biomechanical evaluation rather than heavily depending on the type of metastatic lesion, SINS score or visual radiological appearance alone.

The variability in mechanical response observed in this study is consistent with previous *ex vivo* and computational investigations. Costa et al. (2019) reported that approximately half of the vertebrae with lytic lesions exhibited similar or even higher strength compared to lesion-free controls, suggesting that the impact of metastases on bone mechanics might not be generalised solely based on lesion morphology. Similarly, Palanca et al. (2023) demonstrated that in mechanical testing of metastatic spine segments, some metastatic vertebrae exhibited lower strength than their controls, whereas others were stronger due to reactive bone formation (Mohan et al. 2017). The findings in the present study reinforce this complexity: while vertebrae with lytic lesions generally exhibited lower normalised failure loads (<1), several vertebrae with mixed or blastic lesions demonstrated comparable or superior mechanical performance under compressive loads relative to their controls.

Notably, Specimen ID10 had a large lytic lesion (36% of vertebral volume) in its metastatic vertebra and exhibited a normalised failure load equivalent to that of another spine segment with a blastic lesion in its metastatic vertebra (Specimen ID29). The apparently paradoxical behaviour of Specimen ID10, where the lytic vertebra displayed a higher whole-vertebra failure load than its control in the single-vertebra FE analysis, contrasts with the experimental findings reported by Palanca et al. (2023), in which the same metastatic and control vertebrae failed almost simultaneously and at comparatively low load when tested as part of an intact spine segment. This divergence highlights the fundamentally different mechanical environments encountered in single-vertebra FE models and in segment-level experiments. In the isolated configuration, load transfer is governed primarily by the local trabecular and cortical architecture within the vertebral body, and failure is triggered when a prescribed apparent strain threshold is reached. In contrast, failure load in experimental testing (Palanca et al. 2023), defined as the load for which there was a first sharp decrease of the load in the load-time curve, both vertebrae showed permanent deformation, suggesting similar failure loads for both vertebrae.

It is noteworthy that the spine segment corresponding to Specimen ID10 was the only one classified as osteoporotic according to the 2007 ISCD BMD threshold (<0.08 g cm⁻³) (Engelke et al., 2008), and both metastatic and control vertebrae exhibited nearly identical mean BMD, vertebral height, and broadly comparable cross-sectional geometry. Thus, the unexpectedly high normalised failure load observed for the lytic vertebra in the isolated FE analysis cannot be attributed to global densitometric differences or obvious geometric advantages. Rather, this finding reinforces that lesion topology and its interaction with the surrounding trabecular network can strongly influence how load is redistributed within an isolated vertebral body, even when bulk material properties are similar. Such mechanistic nuances rooted in local rather than global structural differences are not captured by clinical scoring systems like SINS, which do not account for patient-specific load paths or regional variations in mechanical competence.

Quite a few specimens had low failure loads (below 1000N) for both the control and metastatic vertebrae which may indicate that the risk of failure is high in both vertebrae. Hence, a normalised failure load criteria might not be sufficient as a predictive tool for fracture risk and a minimum failure load criterion might be added as a second criterion. In order to confirm this, additional experimental tests on a large number of samples need to be carried out.

The strong correlation between qCT-derived mean BMD and predicted vertebral failure load (R² = 0.74 for metastatic and 0.77 for controls) further supports the use of BMD as a primary determinant of mechanical competence in both healthy and pathological bone. This might be partly explained by the fact that BMD is directly used as an input parameter for material property assignment in the finite element models. These results are in agreement with prior studies (Costa et al. 2019; Wang et al., 2012; Dall’Ara et al., 2012; Crawford et al., 2003) that identified BMD as a robust predictor of vertebral strength. In contrast, the weak correlation between BMC and failure load observed in the present study likely reflects that BMC, being an integrated measure of bone density and bone volume, is insensitive to the structural contribution of mineralised tissue that governs load bearing within the vertebral body. Therefore, although mean BMD is also an averaged parameter and does not capture spatial heterogeneity, the voxel-wise qCT-based material mapping used in this study provides a more spatially resolved and biomechanically meaningful characterisation of bone tissue than bulk, mean densitometric parameters such as BMD or BMC.

The absence of statistically significant differences in failure load or compressive strength between metastatic and control vertebrae should be interpreted with caution. The modest sample size (n = 12 pairs) and variability in lesion morphology, size, and anatomical location limit the statistical power of group comparisons. Moreover, the study used a uniform failure criterion (1.9% apparent strain) originally proposed for osteoporotic bone (Keaveny et al., 2014) in menopausal women, which may not fully capture the complex post-yield behaviour of metastatic tissue. Nonetheless, this choice provides a conservative estimate of strength and ensures comparability with prior SSFE studies.

Analysis of spatial strain fields (Fig. 6-7) revealed that regions of high compressive strain (>0.019) were not only confined to metastatic lesions alone but also occurred near the superior and inferior endplates in both metastatic and control vertebrae. Fourteen out of the twenty-four tested vertebrae showed evidence of endplate involvement, suggesting that endplates act as primary stress-transfer regions prone to high strain concentrations and failure, consistent with earlier experimental observations (Palanca et al. 2023; McKay et al. 2020; Jackman et al. 2016; Fields et al. 2010). Notably, all vertebrae with lytic lesions (Figure 5) exhibited elevated strains at the superior endplate, whereas inferior endplate involvement was more common in vertebrae with mixed lesions (Figure 6). Palanca et al. (2023) similarly reported that approximately 70% of vertebral failures occurred at the superior endplate in their tested spine segments, and that vertebrae with lytic lesions typically failed either within the tissue between the lesion and the endplate or directly adjacent to the endplate. However, in the present study, not all vertebrae with lytic or mixed lesions demonstrated consistent high strain in the superior and inferior endplates, respectively. No such strain localisation near to endplates was, however, detected in the blastic case (Figure 6), likely due to the increased local stiffness and reduced compliance of the sclerotic tissue. The localisation of high-strain initiation around the lytic lesions (Soltani et al. 2024; Palanca et al. 2023), as observed in the strain contour plots, indicates that localised reductions in stiffness might influence early mechanical failure. Conversely, in blastic lesions, peak strain concentrations were found in compliant trabecular zones away from the sclerotic regions (Figure 7), suggesting altered load paths and potential stress concentration at the interface between stiff and compliant tissues. These findings suggest that lesion type modulates local load redistribution (Fereydoonpour et al. 2025) within the vertebral body, with lytic lesions leading to strain amplification in adjacent trabecular regions with further progression towards the endplates, whereas blastic lesions might result in load shielding effects.

In addition to these strain patterns, the spatial localisation of equivalent plastic strain (Figure 8) around the metastatic defects observed in both the large lytic lesion (ID10) and the off-centre blastic lesion (ID29) suggests that asymmetric alterations in local stiffness may introduce an effective bending component even under nominally axial compression. Such stiffness asymmetry, arising from the removal of trabecular bone (lytic) or the presence of a stiff sclerotic mass (blastic), can shift the internal load path away from the vertebral centroid. This produces a combined compression–bending state that concentrates plastic deformation at the lesion–bone interface while leaving the lesion cavity or sclerotic core largely unloaded. Although bending moments were not explicitly applied in the FE model, the observed perilesional plasticity is consistent with this mechanically plausible redistribution mechanism (Fereydoonpour et al. 2025). This interpretation helps explain why vertebrae such as ID10 can exhibit extensive localised plasticity yet still sustain comparatively high apparent failure loads in the isolated-vertebra configuration.

From a clinical perspective, these findings underscore the limitations of categorical scoring systems such as SINS, which may not adequately represent patient-specific variations in mechanical risk (Kwan et al. 2025). Vertebrae receiving similar SINS classifications do not necessarily share similar mechanical behaviour under compressive loads (Figure 9). Lytic lesions tend to weaken vertebrae even when SINS categorises them as stable, whereas mixed lesions may remain mechanically competent despite being radiologically classified as potentially unstable. This decoupling between SINS and mechanical stability supports the integration of qCT-based SSFE modelling to more accurately quantify fracture risk, especially within the clinically challenging transition between the stable (0–6) and potentially unstable (7–12) ranges. Such models could complement current clinical tools by identifying vertebrae that are mechanically vulnerable despite moderate radiological SINS, or conversely, those that remain mechanically competent despite extensive radiological involvement.

However, despite using a robust, reproducible qCT-FE pipeline, this study included a relatively small cohort and lacked direct experimental validation of the predicted mechanical metrics. Accordingly, patient-matched adjacent vertebrae were used as internal controls to support within-subject comparisons. Additionally, under the assumptions that local qCT-derived BMD influence the mechanical behaviour of cancer-affected bone specimens (Stadelmann et al, 2020; Costa et al. 2019; Nazarian et al. 2008; Kaneko et al. 2004), metastatic lesions were treated as homogeneous inclusions within the vertebral body, without explicitly modelling lesion-specific material anisotropy or necrotic regions. Although vertebral bone exhibits anisotropic material properties, the assumption of isotropic bone behaviour included in this study was deemed acceptable given the comparative nature of the study and the focus on relative differences between metastatic and control vertebrae (Ghosh et al. 2025). Moreover, the loading was applied to isolated vertebral bodies, without accounting for the influence of surrounding structures such as intervertebral discs and facet joints, which play an important role in physiological load transfer across the spine (Groenen et al., 2018). The use of a single apparent strain-based failure criterion (Keaveny et al., 2014) across all lesion types may either underestimate or overestimate the true strength when local tissue properties diverge substantially from those of healthy bone. Future studies should investigate anisotropic material behaviour with multi-axial loading, incorporate microstructural information from high-resolution imaging to parameterise lesion-specific material properties and perform direct mechanical validation. Expanding the dataset to include multi-lesion segments and applying machine learning–based surrogate models may further enable clinical translation of this computational approach.

## 5 Conclusions

This study presents a reproducible SSFE modelling pipeline for biomechanical assessment of vertebrae with metastatic lesions, encompassing lytic, blastic, and mixed types. The qCT-based nonlinear heterogeneous models demonstrated that lesion type, size, and surrounding bone adaptation influence vertebral mechanical competence in complex and lesion-specific ways. While lytic lesions generally reduced strength relative to adjacent controls, mixed lesions showed more variable effects, with some cases exhibiting equal or greater strength than lesion-free vertebrae. Across pooled data for this small cohort, no statistically significant differences in strength or failure load were found between metastatic and control groups, underscoring the limitations of using lesion presence or SINS score alone to predict fracture risk. Overall, the proposed SSFE framework provides a robust and scalable methodology for quantitative fracture risk evaluation in vertebrae with metastatic disease. By integrating lesion-specific mechanical insights with patient imaging, this approach lays the groundwork for future clinical decision-support tools aimed at improving personalised treatment strategies in spinal oncology.

## CRediT authorship contribution statement

**Rajdeep Ghosh:** Conceptualisation, Data Curation, Formal Analysis, Investigation, Methodology, Software, Visualisation, Writing – original draft. **Eddie Shearman:** Data Curation. **Robert Roger**: Methodology. **Marco Palanca:** Funding acquisition, Resources, Writing – review & editing. **Enrico Dall’Ara:** Conceptualisation, Funding acquisition, Investigation, Methodology, Project Administration, Resources, Supervision, Writing – review & editing. **Damien Lacroix:** Conceptualisation, Funding acquisition, Investigation, Methodology, Project Administration, Resources, Supervision, Writing – review & editing.

## Acknowledgments

The study was partly funded by the METASTRA project (EU H2022 grant ID 101080135; UK Horizon Europe Guarantee Extension ID: 10075325) and by the Marie Skłodowska-Curie Individual Fellowship (MetaSpine, MSCA-IF-EF-ST, 832430/2018). The funders had no role in the design of the study; in the collection, analyses, or interpretation of data; in the writing of the manuscript; or in the decision to publish the results.

