## Supplementary Materials for "Prediction of compressive strength of vertebral body with metastatic lesions based on quantitative computed tomography-based subject-specific finite element models"

**Supplementary Figure 1**

Supplementary Figure 1 illustrates the European Spine Phantom inserts containing calcium hydroxyapatite (CaHA) with mean reference densities of 0.05 g/cm³, 0.10 g/cm³, and 0.20 g/cm³, covering the full physiological range of cortical and trabecular bone densities observed across different age groups. A densitometric calibration curve (Supplementary Fig. 1c) was generated by calculating the mean Hounsfield Units (HU) within each phantom insert in ImageJ (version 1.54r) and plotting these values against their corresponding reference densities provided by the manufacturer.

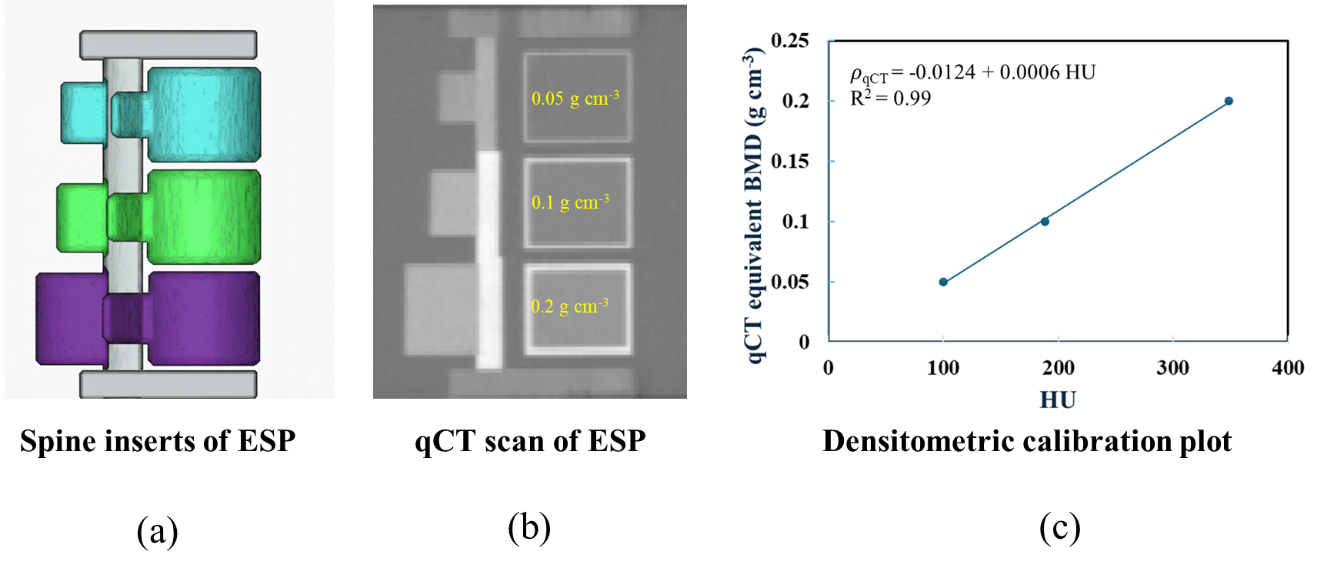

**Supplementary Figure 1.** A representative 3D model of spine inserts of European Spine Phantom (ESP) with three different colours representing inserts with three different densities of CaHa (a); qCT scan of ESP with the imaging parameters mentioned in the text (b); Densitometric-calibration curve with density-BMD relationship (c)
